# An N-terminal ATTR Fibril Segment Promotes Transthyretin Amyloid Nucleation and Polymorphism

**DOI:** 10.64898/2026.09.04.749224

**Authors:** Laxmikant Gadhe, Binh An Nguyen, Maria del Carmen Fernandez-Ramirez, Katerina Konstantoulea, Macy Lozen, Harichandra D. Tagad, Ayde Mendoza-Oliva, Jaime Vaquer-Alicea, Marc I. Diamond, Lorena Saelices, Nikolaos N. Louros

**Affiliations:** Center for Alzheimer’s and Neurodegenerative Diseases, University of Texas Southwestern Medical Center, Dallas, TX, USA; Peter O’Donnell Jr. Brain Institute, University of Texas Southwestern Medical Center, Dallas, TX, 75390, USA; Department of Biophysics, University of Texas Southwestern Medical Center, Dallas, TX, 75390, USA

## Abstract

ATTR amyloidosis is caused by transthyretin (TTR) amyloid deposition, yet the sequence-encoded events linking TTR misfolding to fibril nucleation and structural polymorphism remain incompletely defined. Here, we exploit the modular organization of patient-derived TTR fibrils to investigate two components of the pathological core: an N-terminal β-hairpin spanning residues 11–35 (N-TTR) and a larger C-terminal fragment spanning residues 57–123 (C-TTR). Both fragments independently form β-rich amyloid fibrils, as demonstrated by electron microscopy, circular dichroism, and fluorescence spectroscopy. Yet, their activities differ markedly. N-TTR fibrils promote full-length TTR aggregation and seed in an engineered cellular biosensor platform established to detect templated TTR assembly, whereas C-TTR aggregates show no detectable templating activity. Cryo-electron microscopy reveals two N-TTR polymorphs that preserve structural features of disease-derived folds, while energetic profiling identifies N-TTR as a stabilizing hotspot within ex vivo structures, providing a basis for this templating functionality. These findings reveal a functional hierarchy among amyloidogenic segments of TTR, since distinct regions form fibrils independently, but only those with structural compatibility efficiently template the parent protein. N-TTR therefore represents an autonomous amyloidogenic segment that links local sequence propensity to TTR nucleation, templating, and fibril polymorphism.

## Introduction

Amyloid fibrils are ordered protein assemblies defined by a characteristic cross-β architecture and are associated with a wide range of degenerative and systemic human diseases^1-3^. A defining feature of amyloid formation is that a single precursor protein can give rise to structurally distinct fibril folds, or polymorphs, which may vary across tissues, mutations, disease phenotypes, or biochemical environments^2,4-6^. Understanding how these polymorphs are formed remains a central question in amyloid biology. Although mature fibril cores can encompass large portions of a protein sequence, their assembly is often governed by shorter aggregation-prone regions (APRs)^7-11^ that form intra- or intermolecular β-sheet contacts within protofilament cores and as protofilament cross-interfaces^3,12,13^.

Transthyretin amyloidosis (also referred to as ATTR amyloidosis) is a systemic amyloid disease caused by the deposition of fibrils derived from transthyretin (TTR)^14-16^, a normally soluble tetrameric transport protein^17,18^. TTR deposits can affect multiple organs, particularly the heart and peripheral nerves, and can arise from either wild-type TTR or destabilizing hereditary variants^19-24^. Tetramer dissociation and subsequent monomer misfolding are key upstream events in ATTR pathogenesis^25-27^, but these processes alone do not fully explain how TTR molecules initiate ordered assembly or adopt distinct amyloid folds. Recent cryo-EM studies of patient-derived ATTR fibrils have revealed a growing structural landscape, including fibrils with shared morphologies as well as fibrils displaying patient-, tissue-, or variant-associated polymorphism^20,28-35^. These findings suggest that TTR fibril formation is shaped by both global protein destabilization and local sequence-encoded constraints. A recurring feature of ex vivo ATTR structures is their modular organization. Rather than forming a continuous core encompassing the entire TTR sequence, most ATTR protofilaments contain distinct N-terminal and C-terminal core elements. This organization raises the question of whether the different regions of the ATTR core contribute equally to fibril stability, nucleation, and polymorphism. Prior studies show that local APRs can bias polymorph selection, template aggregation in vitro and in cells, and contribute hierarchically to amyloid assembly^12,36,37^. Patient-derived ATTR structures may therefore provide not only descriptions of mature fibril architecture but also a means of identifying local amyloidogenic segments with functional roles in templating and structural diversification. Whether the distinct regions of the ATTR core possess such autonomous and functionally differentiated assembly properties remains unknown.

Here, we investigated the modular organization of the structural core of ATTR fibrils by evaluating two hemi-fragment constituents: an N-terminal segment spanning residues 11–35, termed N-TTR, and a C-terminal fragment spanning residues 57–123, termed C-TTR. We combined amyloid-propensity prediction, energetic profiling of ex vivo ATTR structures, peptide fibrillization assays, in vitro and cellular seeding experiments, and cryo-EM structure determination. Both fragments formed amyloid fibrils independently, but only N-TTR promoted aggregation of full-length TTR in vitro and in a cellular biosensor. Cryo-EM and energetic analyses further showed that N-TTR forms ATTR-compatible structures and constitutes a stabilizing hotspot within patient-derived fibril folds. These findings suggest a functional hierarchy among amyloidogenic regions of TTR and identify N-TTR as an autonomous self-assembly segment that may connect partial TTR unfolding to fibril nucleation, templating, and polymorphism.

## Results

### Profiling transthyretin aggregation propensity

We first asked whether the modular organization of ex vivo TTR amyloid fibrils reflects the presence of discrete amyloidogenic motifs within the TTR sequence. To this end, we analyzed the TTR sequence using complementary aggregation prediction methods and mapped the resulting scores onto ex vivo ATTR fibril cores. Both CORDAX^38,39^ and TANGO^40^ identified prominent ARPs within TTR, including a strong N-terminal signal spanning the segment corresponding to residues 11–35, as well as APRs within the larger C-terminal core fragment, corresponding to residues 57–127 (**Fig. 1a**). These largely coincide with the hemi-fragments observed in the ex vivo ATTR fibril cores (11-35 and 57-123, hereafter called N-TTR and C-TTR).

**Figure 1.**
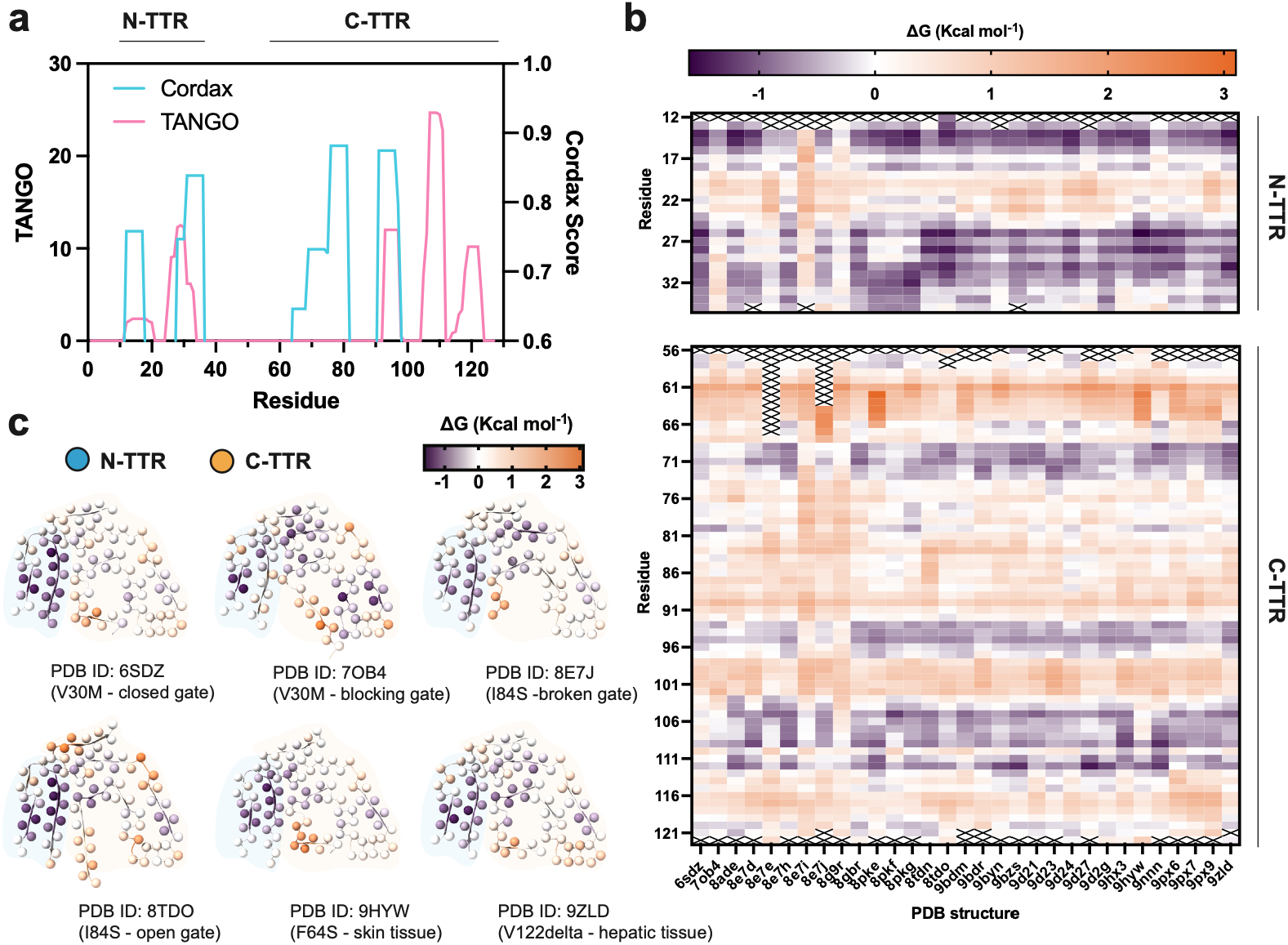
N-TTR is a stabilizing amyloid motif in ATTR fibrils. **(a)** Sequence-based amyloid propensity analysis of full-length TTR using CORDAX and TANGO. The N-terminal TTR segment corresponding to residues 11–35 (N-TTR), and the C-terminal core fragment corresponding to residues 57-123 (C-TTR) are indicated. Both predictors identify amyloid-prone regions within TTR. **(b)** Residue-level energetic profiling of ex vivo ATTR fibril structures. Each row corresponds to an individual TTR PDB structure and each column to a fibril-core residue. Energetic values are shown as ΔG contributions, with stabilizing and destabilizing regions indicated by the color scale. The N-TTR segment is consistently enriched in stabilizing energetic contributions across ex vivo ATTR fibril structures. **(c)** Representative ex vivo ATTR fibril structures colored by per-residue energetic contribution.

To assess whether these amyloidogenic regions also contribute to the stability of disease-derived fibril structures, we next performed energetic profiling at the residue level across available ex vivo ATTR fibril models. This computational analysis estimates the relative energetic contribution of individual residues to the stability of the fibril architecture from their local structural environment and intermolecular contacts^41-44^. Favorable negative energetic contributions therefore identify residues predicted to stabilize the observed fibril conformation, whereas positive values indicate weaker or potentially destabilizing local interactions. Our analysis revealed that the N-TTR region is consistently among the most stabilizing elements across the analyzed ATTR fibril structures (**Fig. 1b**). In contrast, the C-terminal core displayed a more heterogeneous energetic profile, with stabilizing contributions distributed across multiple subregions. Representative ex vivo ATTR fibril structures colored by per-residue energetic contribution further showed that the N-TTR segment forms a recurrent stabilizing β-hairpin within the fibril core (**Fig. 1c**).

### ATTR core fragments form amyloid fibrils in isolation

Guided primarily by the modular organization of the ex vivo TTR fibril core and supported by the aggregation-propensity and energetic analyses above, we synthesized peptides corresponding to its two major structural hemi-fragments: the N-terminal β-hairpin segment, N-TTR (residues 11-35), and the larger C-terminal core fragment, C-TTR (residues 57-123) (**Fig. 2a, S1**). The choice of N-TTR was further supported by a previously proposed pathway for TTR amyloid assembly in which the N-terminal region adopts a hair-pin-like conformation during early oligomerization, suggesting that this segment may have an intrinsic tendency to form an amyloid-competent structural unit^28^. We then tested whether each fragment could form amyloid fibrils independently of the full-length protein. N-TTR assembles rapidly into thioflavin T (ThT)-positive aggregates in phosphate buffer (pH 7.4), (**Fig. 2b**). Similarly, C-TTR formed ThT-positive assemblies, but displayed slower kinetics (**Fig. 2c**). Transmission electron microscopy confirmed that both peptides formed fibrillar assemblies in isolation. N-TTR produced abundant long fibrils, whereas C-TTR formed shorter fibrillar assemblies (**Fig. 2d-e**). Circular dichroism (CD) spectroscopy further supported the acquisition of β-rich secondary structure (**Fig. 2f-g**). Conversion of N-TTR from the soluble state into fibrils was accompanied by a shift in the spectrum from a minimun near 200 nm, typical of a predominantly disordered ensemble, to a minimum near 218 nm, consistent with the presence of β-sheet-rich secondary structure (**Fig. 2f**). C-TTR underwent a similar spectral transition upon fibril formation (**Fig. 2g**). These results show that both hemi-fragments of the ex vivo ATTR fibril core contain sufficient sequence information to assemble into amyloid-like fibrils.

**Figure 2.**
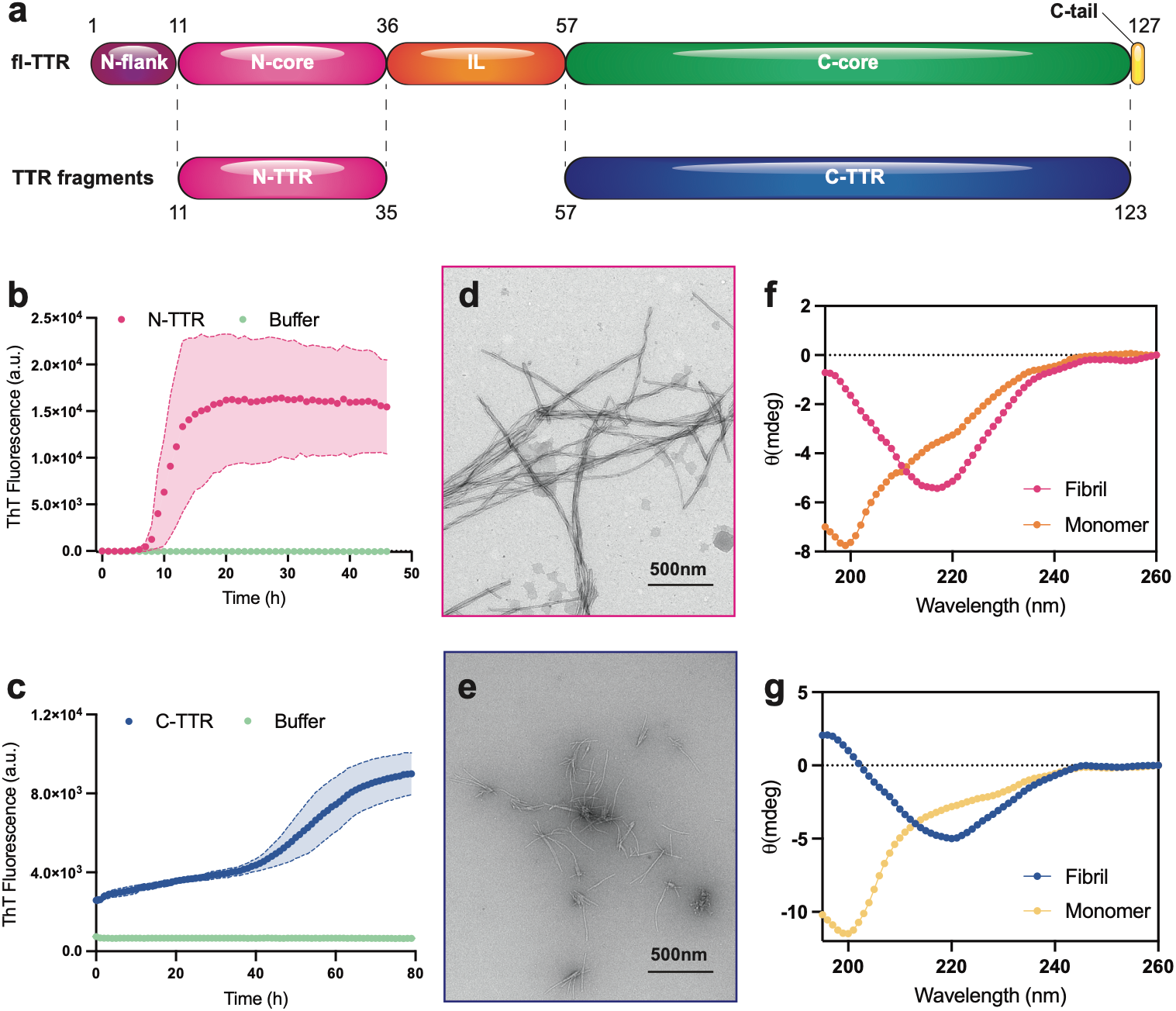
TTR core fragments are intrinsically amyloidogenic. **(a)** Linear schematic of full-length TTR and the synthetic TTR fragments designed in this study. The N-TTR fragment spans residues 11–35, whereas the C-TTR fragment spans residues 57–123. **(b**,**c)** Thioflavin T fluorescence kinetics of isolated N-TTR and C-TTR peptides. **(d**,**e)** Transmission electron microscopy images of N-TTR and C-TTR assemblies. Both peptides form fibrillar structures in isolation. Scale bars, 500 nm. **(f**,**g)** Far-UV CD spectra of monomeric and fibrillar N-TTR and C-TTR.

### N-TTR promotes transthyretin propagation in vitro and in an engineered cellular biosensor

Because both core fragments formed fibrils independently, we next questioned whether either could promote aggregation of full-length TTR. To address this, we used MTTR, an engineered monomeric TTR variant incorporating the F87M/L110M substitutions^45^. Addition of seeds prepared by sonicating preformed N-TTR amyloid fibrils not only markedly accelerates aggregation of MTTR, but also triggers amyloid fibril formation under conditions in which MTTR alone forms amorphous aggregates^46^ (**Fig. 3a, c**). By contrast, C-TTR aggregates show no seeding activity toward MTTR under the same conditions (**Fig. 3b**). Transmission electron microscopy of endpoint samples supports this distinction, as revealing the formation of amyloid fibrils in reactions seeded with N-TTR (**Fig. 3c**). These results indicate that the N-terminal amyloidogenic segment is functionally distinct from the C-terminal segment in its ability to promote aggregation of the parent protein.

**Figure 3.**
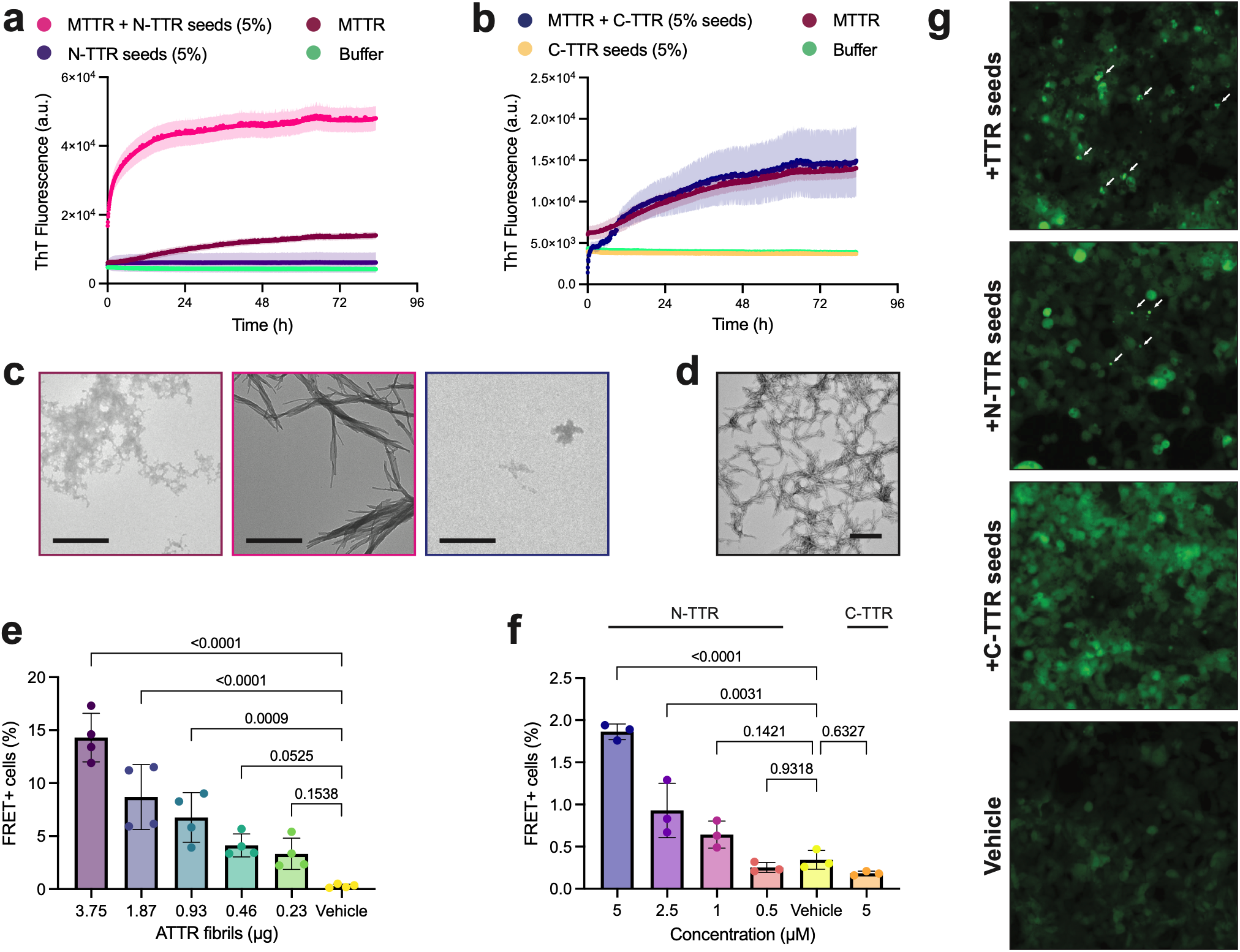
N-TTR selectively seeds TTR aggregation. **(a)** Thioflavin T fluorescence kinetics of full-length MTTR in the absence or presence of preformed N-TTR fibril seeds (n=3 individual repeats). **(b)** Thioflavin T fluorescence kinetics of full-length MTTR in the absence or presence of preformed C-TTR fibril seeds (n=3 individual repeats). **(c)** Electron micrographs of endpoint material derived by incubating MTTR alone (red) or seeded with N-TTR (magenta) and C-TTR preformed aggregates (blue). Scale bar: 1 μm. **(d)** Electron micrograph of amyloid fibrils extracted from cardiac tissue of a patient ATTRwt amyloidosis. Scale bar: 200 nm. **(e)** Cellular TTR biosensor response to ex vivo-derived TTR seeds across the indicated input amounts. TTR seeds induce a dose-dependent increase in FRET-positive cells, validating the sensitivity of the biosensor assay. Statistics: One-way ANOVA with Dunnett’s test for multiple comparisons (n=4 individual repeats). **(f)** Cellular TTR biosensor response to N-TTR and C-TTR seeds. Statistics: One-way ANOVA with Dunnett’s test for multiple comparisons (n=3 individual repeats). **(g)** Representative fluorescence images of TTR biosensor cells treated with TTR seeds, N-TTR seeds, C-TTR seeds, or vehicle. Arrows indicate representative punctate biosensor inclusions.

To investigate whether fibrils formed by the TTR fragments can propagate aggregation of full-length TTR in a cellular environment, we developed a TTR aggregation biosensor cell line following the example of previous models developed for the detection of proteopathic seeding of amyloid proteins^47,48^. In this system, full-length TTR is fused to the photoconvertible fluorescent protein mEOS3.2. Following partial photoconversion from green to red fluorescence, templated aggregation of TTR brings green- and red-labeled molecules into close proximity, generating a FRET signal that provides a sensitive and quantitative readout of intracellular seeding activity. Using this biosensor, we first confirmed robust seeding activity with ATTR fibrils extracted from patient tissue (**Fig 3d**), which induced the highest fraction of FRET-positive cells in a dose-dependent manner (**Fig. 3e**). Although modest, N-TTR fibrils also produced a concentration-dependent increase in FRET-positive cells relative to vehicletreated controls (**Fig. 3f-g**). In contrast, C-TTR fibrils failed to elicit a detectable biosensor response under identical conditions (**Fig. 3f-g**).

Together, these findings demonstrate that, although both N-TTR and C-TTR fragments assemble into amyloid fibrils in vitro, only N-TTR fibrils efficiently template the aggregation of full-length TTR in vitro and in cells.

### Cryo-EM reveals ATTR-compatible N-TTR fibrils

Having established that N-TTR selectively templates TTR aggregation, we next sought to define the structural basis of this activity. We therefore performed cryo-EM analysis of N-TTR fibrils assembled in isolation. Micrographs revealed long, disperse fibrils suitable for helical reconstruction (**Fig. 4a**). Classification identified two major fibril architectures, hereafter referred to as the fast-twisting and slow-twisting N-TTR polymorphs (**Fig. 4b**). Both polymorphs were composed of four laterally associated protofilaments, but they differed in their overall dimensions and helical organization. The fast-twisting polymorph had a fibril width of approximately 8.5 nm, an apparent crossover distance of approximately 50 nm, and a helical rise of 4.77 Å, whereas the slow-twisting polymorph was wider, approximately 11 nm, and showed a longer apparent crossover distance of approximately 170 nm and a helical rise of 4.81 Å (**Fig. 4c**).

**Figure 4.**
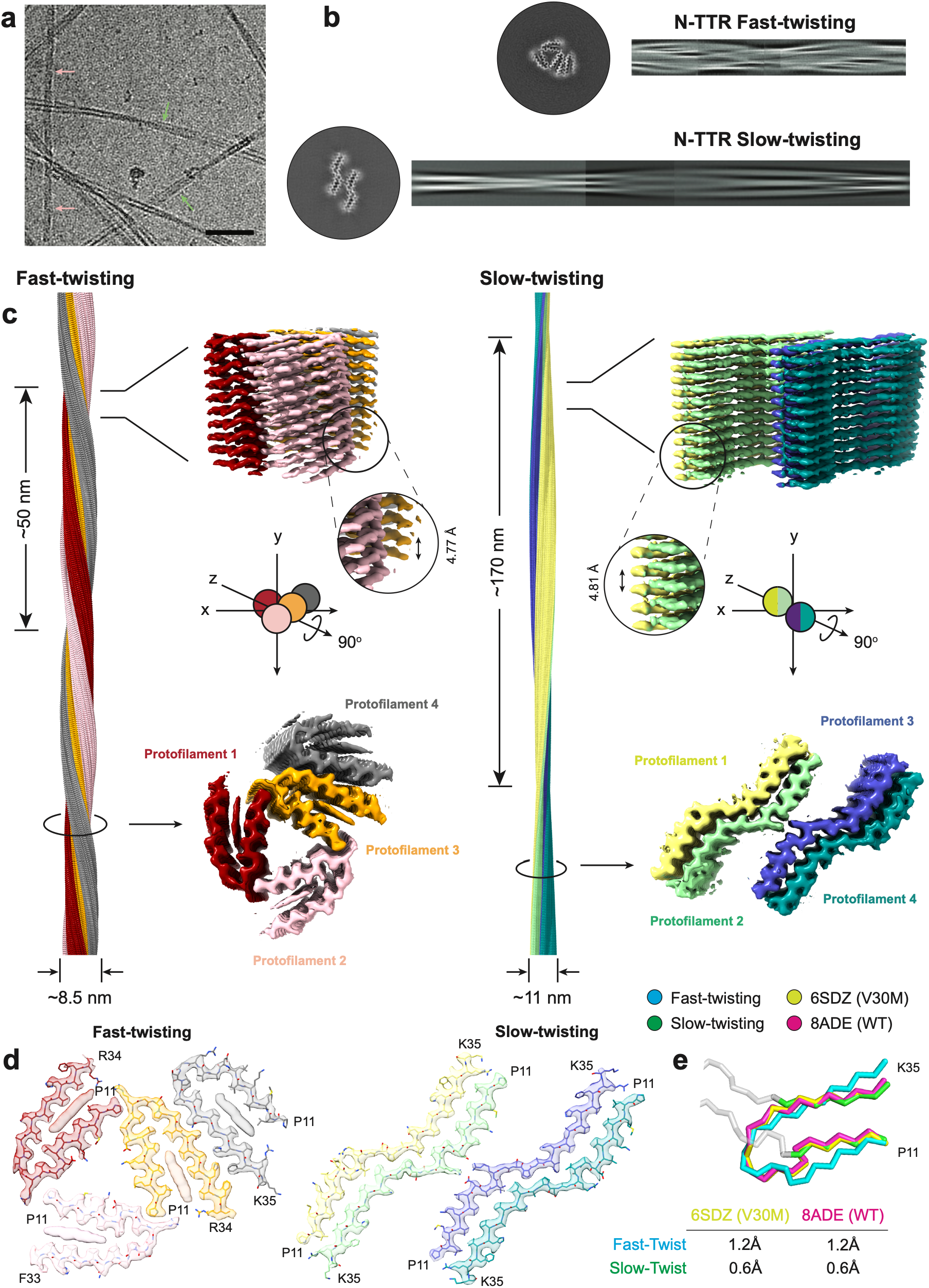
Cryo-EM reveals ATTR-compatible N-TTR fibrils. **(a)** Representative cryo-EM micrograph of N-TTR fibrils formed in isolation. Pink and green arrows show fast and slow twisting filaments. Scale bar: 500 Å. **(b)** Representative 2D class averages and cross-sectional views of two N-TTR fibril morphologies (fast-twist and slow-twist N-TTR). **(c)** Cryo-EM reconstructions and atomic models of the fast-twisted and slow-twisted N-TTR polymorphs. Both fibrils are composed of four protofilaments, shown in distinct colors. **(d)** Cross-sectional views N-TTR fibril polymorphs showing protofilament organization and side-chain packing. **(e)** Structural comparison of N-TTR fibril polymorphs with the corresponding N-terminal segment in ex vivo ATTR fibril structures. Structural alignment RMSD values are shown in the inset table.

The fast-twisting N-TTR polymorph adopted a compact multi-protofilament architecture in which each N-TTR molecule formed a β-hairpin-like amyloid conformation (**Fig. 4c-d**). The four protofilaments packed laterally against one another to generate the fibril core, with residues spanning P11 to K35 defining the ordered segment (**Fig. 4d**). An additional unassigned density was also observed with the fast-twisting protofilament core, but its identity could not be confidently assigned. Notably, this β-hairpin arrangement closely resembles the overall architecture adopted by the corresponding N-terminal segment in ex vivo ATTR fibril cores. Thus, the isolated N-TTR peptide is sufficient to reproduce a disease-associated local fold, supporting the idea that the β-hairpin propensity of residues 11–35 is intrinsically encoded by this amyloidogenic TTR segment.

The slow-twisting N-TTR polymorph adopted a more extended architecture, in which the four protofilaments formed an opened paired assembly held by a continued interface involving all residues of the segment (**Fig. 4c-d**). Despite this distinct protofilament fold, local structural alignment showed that this polymorph also retained side-chain interaction features similar to those observed for the same segment in ex vivo ATTR fibril cores, as residues 11-22 and 24-35 align with the N-terminal fragment (**Fig. 4e**). Indeed, comparison with V30M and WT ex vivo ATTR structures showed that both N-TTR polymorph types preserve ATTR-compatible contact surfaces for the fast-twisting polymorph (alignment of residues 11-35) and the slow-twisting polymorph (alignment over 11-22 and 24-35) (**Fig. 4e**). These findings suggest that the N-TTR segment encodes a preferred amyloid interaction grammar that can be accommodated in more than one fibril state, potentially indicating that the intrinsic structural propensity of this highly amyloidogenic TTR segment may contribute to templating full-length TTR and steering the polymorphic landscape of ATTR fibril formation.

## Discussion

In this work, we identify TTR residues 11–35 as an autonomous amyloidogenic segment that links local sequence propensity to ATTR fibril nucleation and architecture. N-TTR is predicted to be highly amyloidogenic, contributes favorably to the stabilization of ex vivo ATTR fibril structures, forms amyloid fibrils independently of the full-length protein, and selectively promotes TTR aggregation in vitro and in an engineered cellular TTR biosensor. Cryo-EM further shows that the N-TTR fibril architectures reproduce the local conformation found in ex vivo patient-derived ATTR fibrils. This work represents the first recapitulation of part of the ex vivo structure of ATTR fibrils at near-atomic resolution. It also indicates that substantial disease-relevant structural information is intrinsically retained within this relatively short region of the TTR sequence and establishes N-TTR as a tractable reductionist model for future diagnostic and therapeutic development in ATTR.

The contrasting behavior of N-TTR and C-TTR further illustrates that autonomous fibril formation is not sufficient for productive templating. Although C-TTR readily forms β-rich fibrillar assemblies, the most straightforward explanation for its lack of detectable seeding activity is that the isolated C-terminal fibrils are not structurally compatible with the conformation required to recruit the full-length protein. This contrasts with N-TTR, whose isolated fibrils preserve local backbone and side-chain interaction features observed in ex vivo ATTR structures. A similar distinction has emerged in other amyloid systems, including tau, where aggregation propensity alone does not predict templating activity and structurally compatible segments can seed the parent protein more effectively than highly amyloidogenic APRs that adopt alternative packing arrangements^12^. In this context, C-TTR may form off-pathway fibrils that satisfy its intrinsic amyloid propensity without reproducing a productive TTR templating surface. Other, non-mutually exclusive explanations remain possible. Productive C-terminal assembly may require contacts contributed by regions outside the isolated fragment. Sequence compatibility may also contribute: the C-TTR peptide used here corresponds to the regions where the monomeric variant, used for the in vitro and cellular seeding systems, contain mutations that could influence compatibility with a C-terminal templating surface. Finally, C-TTR-mediated seeding may depend on assembly conditions not captured here, including pH, ionic environment, cofactors, proteolytic processing, or fibril maturation state. Regardless of the underlying mechanism, the contrasting behavior of the two fragments establishes that the ability to form amyloid independently does not necessarily confer the ability to template the parent protein.

This distinction is consistent with a broader hierarchy among aggregation-prone regions within amyloid-forming proteins. Short amyloidogenic sequence elements are well established as important determinants of fibril formation because they can form repetitive β-sheet interfaces and steric-zipper-like interactions that stabilize amyloid assemblies^2,3,5,49^. However, sequence-based aggregation propensity alone does not identify which regions are capable of productive nucleation in the context of the intact protein. N-TTR satisfies a broader set of criteria: it is strongly amyloidogenic by sequence, contributes favorably to the energetic stabilization of disease-derived fibril cores, forms ordered fibrils in isolation, promotes aggregation of full-length TTR, and retains ATTR-compatible structural features. These observations support the idea that amyloid-forming proteins may contain multiple amyloidogenic segments with distinct functional roles, only a subset of which are sufficiently compatible with the structural constraints of the parent protein to nucleate or propagate its assembly.

The structural basis for this functional modularity may lie in the defined compact β-hairpin fold of N-TTR. Unlike C-TTR, which corresponds to a more spatially dispersed region of the ex vivo fibril core, N-TTR forms a self-complementary contact network. This intramolecular closure allows a single 25-residue stretch to establish a complete, self-complementary steric-zipper interface without requiring long-range contacts. By contrast, C-TTR may depend on stabilizing contacts supplied only within the complete full-length fold. This interpretation is consistent with structural studies demonstrating that other amyloid-forming proteins often adopt β-hairpin conformations in isolation^50-54^, and with efforts to direct amyloid assembly and end-state folds through synthetic hairpin-like derivatives^55-61^. Previous studies of shorter peptides further support this. In particular, a short APR derived from strand B (residues 28-33) forms fibrils with an unusual out-of-register steric zipper architecture^62^ proposed in several amyloid systems to represent potentially assembly-competent or toxic oligomeric states^63,64^.

This behavior is also consistent with the concept of framework polymorphism, in which conserved amyloidogenic segments are often re-used and alternatively embedded within different polymorphic folds^36,43^. In this model, N-TTR provides a local stabilizing amyloidogenic segment whose preferred interaction grammar can be retained even as the surrounding fibril architecture changes^5,13,65^. This concept tracks with an emerging view of amyloid fibrils as modular structures in which local sequence segments structurally bias the accessible conformational landscape, potentially steering amyloid templating^12,59,61^. Thus, polymorph-guiding amyloidogenic segments may represent a general principle across amyloid systems: local motifs can act as structural anchors that constrain assembly while still permitting overall polymorphic variation. This framework potentially explains why the protofilament fold variability of ATTR fibrils from peripheral tissues is detected solely in the C-terminal segment of the core and may help reconcile recent ATTR structural studies that report structures of TTR in the brain. Both fibrils isolated from the brains of patients carrying the V30M and V30G variants differ substantially from those reported previously from peripheral tissues^66^. Within these structures, the N-terminal segment is the only structurally preserved segment in V30M, with the C-terminal part wrapping around it in a different final conformation, but is missing in the reduced ordered V30G core, potentially due to proteolytic processing. Thus, TTR polymorphism may be constrained by local energetics of such amyloidogenic segments rather than generated by entirely unrelated packing solutions.

The autonomous behavior of N-TTR also provides a plausible molecular link between partial unfolding of native TTR and amyloid nucleation. Previous studies suggest that TTR misfolding involves displacement of the peripheral β-strands and their connecting loop^67-69^. In the native fold, experimental evidence highlights the local destabilization and unfolding of C and D β-strands^69-71^. These elements shield or conformationally restrain the A–B facing β-strands encompassing residues 11–35, and their displacement could therefore expose N-TTR, enabling this otherwise protected segment to promote nucleation. Such a mechanism would be consistent with the observation that partial unfolding promotes TTR aggregation.

This mechanism also provides a potential therapeutic angle downstream of native tetramer stabilization. Current ATTR therapies primarily act by reducing TTR production or stabilizing the native tetramer^72-76^. The identification of a discrete initiator within residues 11–35 suggests that amyloid assembly itself may provide additional intervention points. Because N-TTR is energetically favored within ex vivo fibril cores, can promote full-length TTR aggregation, and reproduces disease-compatible structural contacts in isolation, its interaction surfaces could provide templates for the development of conformation-selective binders, fibril-capping molecules, or inhibitors that interfere with productive intermolecular association^60,77-82^. Such approaches could complement strategies directed at the native precursor by targeting structural events that occur after TTR destabilization.

## Materials and Methods

### Ethical statement

Ex vivo ATTR fibrils were extracted from fresh-frozen left ventricular cardiac tissue obtained postmortem from an 84-year-old male patient with ATTRwt amyloidosis (ATTRwt-p3), whose amyloid fibril structure had been previously determined by cryo-electron microscopy^83^. The sample was originally provided by the laboratory of the late Dr. Merrill D. Benson at Indiana University. The Office of the Human Research Protection Program granted expedited approval from the Internal Review Board review because the specimen was anonymized. All applicable ethical regulations for research were followed.

### Computational predictions and analysis

Calculations of aggregation propensity were performed using aggregation predictors TANGO^40^ and CORDAX^38,39^. The former being a sequence-based predictor that relies on hydrophobicity and β-sheet propensity to identify aggregation-prone regions, whereas CORDAX is a machine-learning structure-based predictor that reveals compatibility with the amyloid cross-β architecture. Profiling fibril stabilities was performed by applying a thermodynamic profiling framework^41-44^ to ex vivo ATTR structures. Local structural alignment was performed using the Matchmaker function in ChimeraX (v1.10.1). Structural analysis and molecular graphics were generated using ChimeraX (v1.10.1) and statistical analysis and plots were generated using Graphpad Prism (v11.0.2).

### Protein purification

Engineered monomeric transthyretin (MTTR) variant carrying the F87M and L110M substitutions was expressed in *Escherichia coli* Rosetta™ 2 (DE3) competent cells (Sigma-Aldrich). Bacterial cultures were grown in LB medium supplemented with 50 µg/mL kanamycin at 37 °C with orbital shaking at 280 rpm until reaching an optical density at 595 nm of 0.4–1.0. Protein expression was induced by the addition of isopropyl β-D-1-thiogalactopyranoside (IPTG) to a final concentration of 1 mM, and cultures were incubated for an additional 4 h under the same conditions. Cells were harvested by centrifugation at 6,000 × *g* for 20 min. Cell pellets were resuspended in 30 mL of affinity binding buffer per liter of bacterial culture (20 mM Tris-HCl, 300 mM NaCl, 20 mM imidazole, pH 7.5), supplemented with two Complete™ Mini EDTA-free Protease Inhibitor Cocktail tablets (Roche; cat. no. 11836170001). Cells were lysed by probe sonication for a total sonication time of 10 min using alternating ON/OFF pulses at 30–40% amplitude. Cell lysates were clarified by centrifugation at 14,000 × *g* for 45 min, and the resulting supernatant was filtered through a 0.45 µm pore-size membrane. Recombinant MTTR was purified using an ÄKTA pure™ chromatography system (Cytiva). The clarified lysate was loaded onto a HisTrap™ HP column (Cytiva), and bound protein was eluted using a buffer containing 20 mM Tris-HCl, 300 mM NaCl, and 500 mM imidazole, pH 7.5. MTTR was further purified by size-exclusion chromatography using a Hi-Load™ 16/600 Superdex™ 75 pg column (Cytiva). Pure protein fractions were pooled and concentrated using Amicon® Ultra centrifugal filter units (10 kDa MWCO; Millipore; cat. no. UFC901008). Protein concentration was determined by measuring the absorbance at 280 nm using a NanoDrop spectrophotometer and the theoretical molar extinction coefficient.

### Fibril extraction and validation

*Ex vivo* ATTR fibrils were extracted from the cardiac tissue of a patient with ATTRwt amyloidosis, whose amyloid fibril structure had been previously determined by cryo-electron microscopy^83^. Fibrils were extracted following a previously described protocol^28,84^ with minor modifications. Briefly, approximately 200 mg of fresh-frozen tissue was thawed, finely minced with a scalpel, and resuspended in 1 mL of Tris-calcium buffer (20 mM Tris, 150 mM NaCl, 2 mM CaCl_2_, 0.1% NaN_3_, pH 8.0). The sample was centrifuged for 5 min, the supernatant was discarded, and the pellet was resuspended in Tris-calcium buffer. This washing step was repeated four times. The washed pellet was then resuspended in 1 mL of Tris-calcium buffer supplemented with 5 mg/mL collagenase and incubated overnight at 37 °C with orbital shaking at 400 rpm. Following centrifugation at 3,100 × *g* for 30 min at 4 °C, the resulting pellet was resuspended in 1 mL of Tris-EDTA buffer (20 mM Tris, 140 mM NaCl, 10 mM EDTA, 0.1% NaN_3_, pH 8.0) and centrifuged at 3,100 × *g* for 5 min. The supernatant was discarded, and the pellet was resuspended in fresh Tris-EDTA buffer prior to the next 5 min centrifugation. This washing step was repeated ten times. Amyloid fibrils were subsequently eluted by gently resuspending the pellet in 200 µL of 5 mM EDTA prepared in ultrapure water, followed by centrifugation and collection of the supernatant. Six sequential elutions were collected from the sample. Except for the overnight collagenase digestion, all extraction steps and buffers were performed and maintained at 4 °C. All sequential elutions were examined using transmission electron microscopy (TEM), and only those containing the highest abundance of amyloid fibrils with minimal tissue debris and other contaminants were pooled. Protein concentration was determined using the bicinchoninic acid (BCA) assay, and the final fibril preparation was adjusted to 0.5 µg/µL.

### Peptide synthesis

Peptides corresponding to the N-TTR (11-35) and C-TTR (57-123) peptide cores of TTR protein were synthesized using Fmoc solid-phase peptide synthesis on a MultiPep 2 robot peptide synthesizer (CEM). Peptides were cleaved, ether-precipitated, and stored at –20 °C. Both peptides were purified using HPLC (Shimadzu Nexera), and purity (>99%) was confirmed by LC–MS (Shimadzu LCMS-2050). The lyophilized peptides were dissolved in 1,1,1,3,3,3-Hexafluoro-2-propanol (HFIP). HFIP was evaporated under nitrogen flow, and the resulting films were reconstituted in the appropriate buffer for downstream experiments.

### Fluorescence assays

Peptide samples (1 mg/mL) were dissolved in 20mM phosphate buffer (pH 7.4) and incubated in a 96-well clear bottom plate at 37 °C under orbital shaking at 300 rpm for 30 sec before each reading using a FLUOstar Omega plate reader (BMG Labtech), using a 448 nm excitation and 482 nm emission wavelength. Samples were mixed with 25 µM ThT in 96-well plates (Greiner #675096). For seeded reactions, 0.5 mg/ml of MTTR dissolved in 10mM sodium acetate (pH 4.3) buffer, including 100 mM KCl, was incubated at 37°C under orbital shaking at 300rpm before each reading using a CLARIOstar plate reader (BMG Labtech) in the presence and absence of preformed N-TTR and C-TTR seeds, generated by sonication of mature fibrils at 20% amplitude using a 3/1s ON/OFF pulse cycle for a total sonication time of 3 min. Data analysis was performed with Mars Omega (BMG Labtech).

### Circular dichroism spectroscopy

Far-UV CD measurements were performed on peptide samples diluted in 20mM phosphate buffer (pH 7.4) at a concentration of 40 µM using a J-815 CD spectrometer with a path length set to 0.1 cm quartz cell at room temperature. The graph was plotted using GraphPad Prism (v10.5.0).

### Generation and maintenance of TTR biosensor cells

TTR biosensor cells were generated in a HEK293T background using the lentiviral expression strategy previously described for tau biosensor cells^85^. Briefly, the cassette region of the FM5-CMV-nk-cassette-linker-FP backbone was replaced with a sequence encoding mature human TTR lacking its N-terminal signal peptide and containing the amyloidosis-associated I84S substitution together with the monomerizing F87M and L110M substitutions. The resulting ΔSP-hTTR(I84S/F87M/L110M) sequence was fused to C-terminal mEos3.2 through the previously described 12-amino-acid linker (GSAGSAAGSGEF). Lentivirus generated from this construct was used to transduce HEK293T cells and establish a stable biosensor cell line. Under basal conditions, TTR-mEos3.2 fluorescence was diffusely distributed throughout the cytoplasm and nucleus, whereas exposure to TTR seed-containing samples induced the formation of intracellular fluorescent inclusions. HEK293T biosensor cells expressing I84S/F87M/L110M TTR-mEos3.2 were maintained in 10 cm culture dishes in Dulbecco’s modified Eagle’s medium (DMEM; Gibco) supplemented with 10% fetal bovine serum (HyClone), 1% penicillin–streptomycin (Gibco), and 1% GlutaMAX (Gibco). Cells were cultured at 37°C with 5% CO_2_ and ≥80% relative humidity. Prior to the experiments, cells were tested for mycoplasma contamination and confirmed to be negative.

### Cellular seeding assays

Biosensor cells were plated in 96-well plates at 25,000 cells per well in 120 µL of complete medium. After 18 h, cells were treated with N-TTR and C-TTR fibrils at final concentrations of 0.5–5 µM and 5 µM, respectively, or with wild-type TTR *ex vivo* fibrils extracted from human cardiac tissue at 0.23–3.75 µg per well. 1× Dulbecco’s phosphate-buffered saline (1× DPBS; Gibco) was used as a vehicle control. All samples were sonicated before use at 65% amplitude for 5 min using 30-s on/off cycles with a QSonica sonicator. For each well, liposome complexes were prepared in 30 µL containing 14.17 µL of Opti-MEM (Gibco), 0.83 µL of Lipofectamine 2000 (Invitrogen), and 15 µL of 1× DPBS containing the indicated sample. Complexes were incubated for 30 min at room temperature and added to the cells, resulting in a final volume of 150 µL per well. Cells were incubated with the complexes for 48 h at 37°C. Cells were harvested with 0.05% trypsin (Gibco) for 5 min at 37°C, quenched with fresh medium, and transferred to 96-well U-bottom plates. After centrifugation, cells were fixed in 2% paraformaldehyde for 10 min and resuspended in 150 µL of cold 1× DPBS.

### Flow cytometry

Fixed cells were partially photoconverted under UV light for 30 min using a Chauvet LED Shadow light source. Cells were then analyzed on an Attune CytPix flow cytometer equipped with a CytKick autosampler (Thermo Fisher Scientific). Green mEos3.2 fluorescence was detected using 488 nm excitation and a 530/30 nm filter, whereas photoconverted red mEos3.2 fluorescence was detected using 561 nm excitation and a 620/15 nm filter. FRET was measured by exciting the donor at 488 nm and detecting emission through a 695/40 nm filter in the PerCP channel. Cells were gated based on forward scatter (FSC) and side scatter (SSC), followed by singlet selection and identification of green- and red-double-positive cells, within which FRET was quantified. The FRET gate was adjusted in Attune Cytometric Software v7.1 using lipofectamine-only samples as negative controls and cells treated with *ex vivo* ATTR fibrils as positive controls. The same gating boundaries were applied across all conditions and experimental replicates. FRET was reported as the percentage of FRET-positive cells per replicate. Data were analyzed using FlowJo v10 (Tree Star). N-TTR and C-TTR conditions were analyzed in triplicate, and *ex vivo* ATTR fibril conditions were analyzed in quadruplicate.

### Transmission electron microscopy

Suspensions (5 μL) of fibril samples were loaded onto glow-discharged carbon-coated Formvar EM grids (Electron Microscopy Sciences) and incubated for 5 min. The grids were washed with Milli-Q water and stained with 2% w/v uranyl acetate for 2 min. The excess stain was removed by blotting with filter paper. The grids were air-dried, and imaging was performed using a transmission electron microscope (JEOL JEM-1400) at an accelerating voltage of 120kV.

### Cryo-EM grid preparation, screening, and data collection

Suspensions (3.5μL) of N-TTR amyloid filaments, prepared in Milli-Q, were loaded onto a glow-discharged holey carbon film grid (Quantifoil R 1.2/1.3, Cu 300 mesh), blotted for 3 sec, and vitrified in liquid ethane using a Vitrobot Mark IV (Thermo Fischer Scientific). Grids were screened on a 200 kV Talos Arctica, and final datasets were collected on a 300 kV Titan Krios microscope equipped with K3 detector (Thermo Fisher Scientific) at the Cryo-Electron Microscopy Facility at The University of Texas South-western Medical Center. Pixel size, frame rate, dose rate, final dose, and number of micrographs per sample are provided in Table S1.

### Helical reconstruction

The raw movie frames were gain-corrected, aligned, motion corrected, and dose-weighted using the motion correction implementation in RELION 5.0^86,87^. Contrast transfer function (CTF) parameters were estimated using CTFFIND 4.1^88^. All subsequence helical reconstruction including two-dimensional classification, three-dimensional (3D) refinement, and post-process were carried out in RELION 5.0. The filaments were picked automatically using Topaz in RELION 5.0. Particles were extracted using a box size of 256 pixels (type I fibrils) and 448 pixels (type II fibrils) with an inter-box distance of 3 and 8 asymmetrical units at helical rise of 4.75 Å, respectively. 2D classifications were used to remove suboptimal segments and the high-resolution classes showing clear fibril layers separation were stitched together to estimate the crossover distance. Initial 3D reference models were generated from a subset of 2D class averages using *relion_helix_inimodel2d* script^89^. Fibril helix was assumed left-handed for 3D reconstruction. Multiple rounds of 3D classifications were performed and particles yielding the highest map quality were selected for iterative 3D auto refinements with optimization of helical twist and rise once estimated resolution of the map reached beyond 4.75 Å. An additional round of 3D classification without image alignment was applied to further remove remaining suboptimal classes. Sequential rounds of Bayesian polishing^90^, CTF refinements^91^ and 3D auto refinements were performed to maximize resolution^92^. Throughout reconstruction, C1 symmetry was applied. Overall resolutions were estimated from Fourier shell correlations at 0.143 between two independently refined half-maps using a soft-edged solvent mask^89^ (Fig. S2). Local resolutions were estimated using the RELION local resolution tool. Additional processing details are available in Supplementary Table 1.

### Model building and refinement

We used the automated machine-learning ModelAngelo approach with minor modifications to obtain an initial atomic model^91^. First, map density was trimmed to a single fibril layer in UCSF ChimeraX (v1.8) ^93^, and the resulting map was submitted to ModelAngelo with the N-terminal TTR sequence ranging residues 11-35 to generate the initial atomic models. Using COOT (v0.9.8.1), we made residue modifications and real-space refinements to finalize the model. Further refinement was carried out using ‘phenix.real_space_refine’ from PHENIX (v1.20)^94^. ChimeraX (v1.8) was used for molecular graphics and structural analysis^93^. Model statistics are summarized in Supplementary Table 1.

## Acknowledgements

We thank the individuals and their families for donating tissue for research and acknowledge the Indiana University Brain Bank for making these valuable samples available. NL and the Louros laboratory were supported by a scholarship from the Thomas O. Hicks Scholar in Medical Research. KK was partially supported by the O’Donnell Brain Institute (OBI) Sprouts Grant Program. LS and the Saelices laboratory were supported by the National Institutes of Health, National Heart, Lung, and Blood Institute, the UTSW Milestone Award, and AstraZeneca. We acknowledge the assistance of the UT Southwestern Cryo-Electron Microscopy Facility (CEMF), Electron Microscopy Core Facility (EMCF), and Structural Biology Laboratory (SBL) for technical support and access to instrumentation. The SBL and CEMF are partially supported by the Cancer Prevention & Research Institute of Texas (CPRIT; grant RP220582), and the EMCF is supported by NIH grant 1S10OD021685-01A1. Computational analyses were performed using the BioHPC high-performance computing facility supported by the Lyda Hill Department of Bioinformatics at UT Southwestern Medical Center. The views expressed in this work are those of the authors and do not necessarily reflect the official policies of the National Institutes of Health.

## Author contributions

NL conceived and initiated the study. LG, MFR, BAN, ML, JVA, AMO, and KK performed the experiments. KK and HDT performed peptide synthesis. NL performed the computational analyses. LG, BAN, JVA, MFR, AMO, MID, LS, and NL analyzed and interpreted the data. MID, LS and NL acquired funding and provided resources. LS and NL supervised the project. LG, BAN, LS, and NL prepared the manuscript, and all authors reviewed and edited the final version.

## Data Availability statement

All experimental and computational raw data generated are available from the corresponding author on reasonable request.

