## Supplementary Information for "An N-terminal ATTR Fibril Segment Promotes Transthyretin Amyloid Nucleation and Polymorphism"

**Supplementary Table 1.** Data collection and refinement parameters

| | | |
|---|---|---|
| Data collection | Type I (Curvy) | Type II (Straight) |
| Microscope | Titan Krios | |
| Acceleration Voltage (kV) | 300 | |
| Detector | K3 | |
| Software | SerialEM 3.5 | |
| Magnification | 105,000x | |
| Pixel size at detector (Å/px) | 0.413 | |
| Defocus range (µm) | -2.4 to -1.0 | |
| Total dose (e/Å$^2$) | 50 | |
| Exposure time (sec) | 2.2 | |
| Number of movie frames | 50 | |
| Usable micrographs | 6475 | |
| Box size (pixel) | 256 | 448 |
| Total extracted segments | 782,992 | 4,126,980 |
| Number of segments after 2D | 228,477 | 3,281,481 |
| Number of straight segments | n/a | n/a |
| Number of segments after 3D | 41,603 | 118,084 |
| Symmetry imposed | C1 | C1 |
| Helical rise (Å) | 4.77 | 4.81 |
| Helical twist (° ) | -1.73 | -0.448 |
| Crossover length (Å) | 496 | 1,932 |
| B factor | -69.5 | -95.9 |
| Map resolution (Å; FSC=0.143) | 3.1 | 3.3 |
| Map resolution (Å; FSC=0.5) | 4.2 | 4.3 |
| Non-hydrogen atoms | 3610 | 3800 |
| Protein residues | 480 | 500 |
| Number of chains | 20 | 20 |
| Water/ligands | 0 | 0 |
| MolProbity score | 1.77 | 1.67 |

| | | |
|---|---|---|
| Clash score | 8.85 | 8.60 |
| Rotamer outliers (%) | 0.00 | 0 |
| R.M.S deviations bonds (Å) | 0.003 | 0.004 |
| R.M.S deviation angle (° ) | 0.546 | 0.630 |
| Ramachandran Plot<br>Favored<br>Allowed<br>Outliers | <br>95.68<br>4.32<br>0.00 | <br>96.74<br>3.26<br>0.00 |
| CaBLAM outliers (%) | 0.5 | 1.19 |
| Model vs Data | 0.8 | 0.81 |

**Supplementary Figure 1.** Analytical HPLC traces of synthetic N-TTR and C-TTR peptides. Peak retention times and relative peak areas are shown in the inset table.

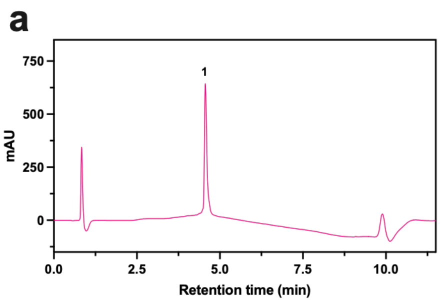


| | Peak # | Ret. time (m) | Area (%) |
|---|---|---|---|
| N-TTR | 1 | 4.56 | 100 |
| C-TTR | 1 | 4.62 | 99.1 |
| | 2 | 6.02 | 0.9 |

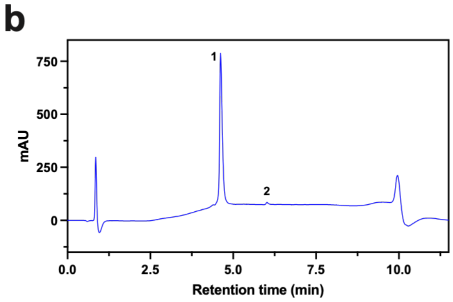

**Supplementary Figure 2. Fourier shell correlation analysis of the N-TTR cryo-EM reconstructions.** Fourier shell correlation (FSC) curves for the **(a)** twisted and **(b)** straight N-TTR fibril reconstructions. Curves show the corrected FSC (black), FSC calculated from the unmasked half-maps (green), FSC calculated after masking (blue), and FSC calculated after phase randomization of the masked maps (red). The corrected FSC accounts for correlations introduced by masking. Map resolution was estimated from the corrected FSC using the FSC = 0.143 criterion.

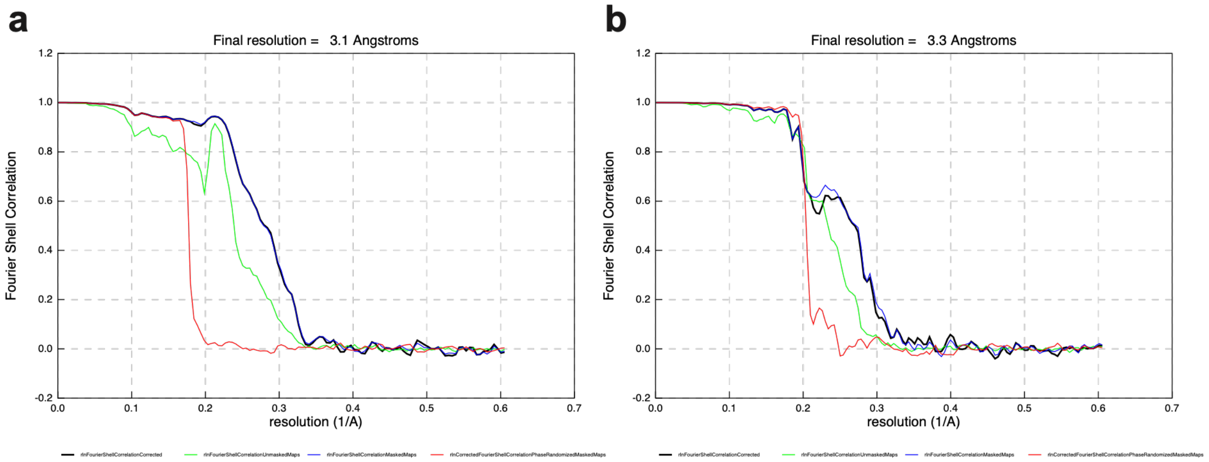
